# Deep conservation of ribosome stall sites across RNA processing genes

**DOI:** 10.1101/2020.09.17.301754

**Authors:** Katarzyna Chyżyńska, Kornel Labun, Carl Jones, Sushma N. Grellscheid, Eivind Valen

**Affiliations:** Computational Biology Unit, Department of Informatics, University of Bergen, Bergen 5020, Norway; Department of Biological Sciences, Durham University, DH1 3LE, UK; Department of Biological Sciences, University of Bergen, Bergen 5020, Norway; Sars International Centre for Marine Molecular Biology, University of Bergen, Bergen 5008, Norway

## Abstract

The rate of translation can vary considerably depending on the mRNA template. During the elongation phase the ribosome can transiently pause or permanently stall. A pause can provide the nascent protein with the required time to fold or be transported, while stalling can serve as quality control and trigger degradation of aberrant mRNA and peptide. Ribosome profiling has allowed for the genome-wide detection of such pause and stall sites, but due to library-specific biases, these predictions are often unreliable.

Here, we address this by taking advantage of the deep conservation of the protein synthesis machinery, hypothesizing that similar conservation could exist for functionally important positions of ribosome slowdown - here collectively called stall sites. We analyze multiple ribosome profiling datasets from a phylogenetically diverse group of eukaryotes: yeast, fruit fly, zebrafish, mouse, and human and identify conserved stall sites. We find thousands of stall sites across multiple species, with proline, glycine, and negatively charged amino acids being the main facilitators of stalling. Many of the sites are found in RNA processing genes, suggesting that stalling might have a conserved regulatory effect on RNA metabolism. In summary, our results provide a rich resource for the study of conserved stalling and indicate possible roles of stalling in gene regulation.

## Introduction

Besides encoding the amino acid sequence of a protein, the coding region of mRNA can contain secondary signals affecting the regulation of the gene. These regulatory signals can modulate elongation rates, leading to translation bursts and pauses and determine how efficiently proteins are synthesized [48, 52]. While many of these signals are likely tuning the rate of synthesis in a subtle fashion, some signals have been shown to cause longer-lasting pauses, or *‘*stalls*’*.

These stalls can have important biological consequences allowing time for the recruitment of various machinery to facilitate subsequent processes, such as membrane targeting or co-translational protein folding [17, 22, 52]. For instance, pausing upon the emergence of the signal peptide from the ribosome exit tunnel promotes recruitment of the signal recognition particle and subsequent targeting of secretory proteins to the endoplasmic reticulum [44]. Slowing down translation downstream of protein structural domains would in turn allow time for the domains to fold into lower-energy folding intermediates [45, 52]. However, if the ribosome stalls due to aberrant translation, it may trigger recruitment of ribosome-associated protein quality control machinery to degrade the nascent peptide through pathways such as nonsense-mediated decay (NMD) or no-go decay (NGD) [9, 17, 31, 51].

Several causes for stalling have been suggested, such as 1) specific amino acids (e.g. proline) in the P- and A-site attenuating the rates of peptide bond formation [3, 49], 2) positively charged residues [14] or non-optimal codon clusters in the nascent peptide, interacting with ribosome exit tunnel [30, 44] or 3) mRNA secondary structure blocking progression of translating ribosomes. [60, 23]. However, it is unclear how widespread each of these causes are and to what extent these are functional.

Ribosome profiling, the sequencing-based transcriptome-wide capture of ribosomal occupancy, can offer unique insight into the dynamics of ribosome translocation based on the distribution of sequencing reads from ribosome protected fragments. Indeed, previous analyses of ribosome profiling data have revealed widespread presence of sites with a high abundance of reads, assumed to be strong ribosomal pauses [29]. However, past studies have also demonstrated a wide range of biases in ribosome profiling data as well as high local variations between data from individual libraries and experiments [3, 5, 24, 26]. Analysis of a single library or experiment is therefore particularly vulnerable to biased results which again can confound the search for regulatory mechanisms.

Here, we address this by taking advantage of the deep conservation of translation machinery suggesting that this may also imply similar mechanisms for stalling. We collect 20 publicly available ribosome profiling datasets for five phylogenetically diverse model organisms in order to identify stall sites conserved across multiple species and experiments. By their conservation, these stalls are likely to represent functional sites. We characterize their biological contexts and uncover signs of conserved mechanisms as well as a conserved role for stalling in RNA processing.

## Results

To identify stall sites that are conserved across species and that are robust to library preparation we collected publicly available ribosome profiling libraries from yeast (*Saccharomyces cerevisiae*), fruit fly (*Drosophila melanogaster*), zebrafish (*Danio rerio*), mouse (*Mus musculus*) and human (*Homo sapiens*). The choice of libraries was dictated by availability, sufficient level of coverage and different experimental conditions, to be able to eliminate noise originating from particular experimental protocol or sequencing bias. All of the data are from wild type and unstressed cells.

After mapping the ribosomal footprints to the transcriptome (see Methods) we identified, for each ribosome protected fragment, the codon currently under translation (P-site). Importantly, this is a library-specific process and requires offsetting each fragment length by a specific amount calculated using Shoelaces [8] (Supplementary Table 1). This processing resulted in codon-resolution data as revealed by ribosome meta profiles (Figure 1A and Supplementary Figure 1), showing clear peaks over the first and/or last translating codons, increased coverage within the coding region (CDS), and in most cases three-nucleotide periodicity, as expected of a correctly P-shifted data [28, 8].

**Fig. 1:**
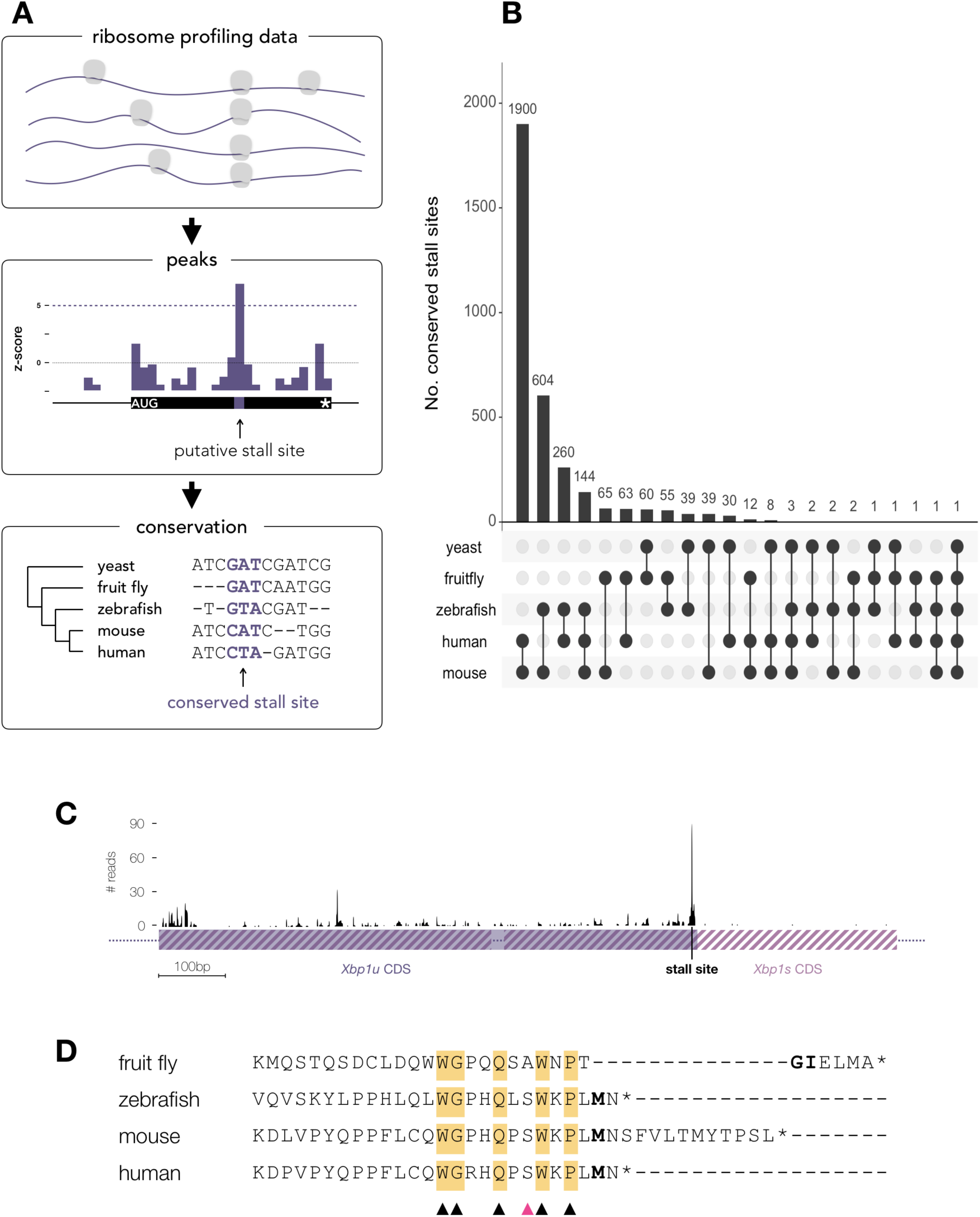
Conservation of stall sites across divergent eukaryotes. (**A**) Schematic representation of stall site analysis. (**B**) Number of conserved stall sites in homologous genes in yeast, fruit fly, zebrafish, mouse and human. Some stall sites are common in lower and higher vertebrates, indicating their importance in translation regulation. (**C**) Ribosome profile on the *Xbp1* mRNA (here values for H3 library). The schematic of two isoforms is shown, the unspliced, shorter *Xbp1u* (solid purple) and spliced, with 3*’* extension, *Xbp1s* (striped purple). The stall site at the 3*’*end of *Xbp1u* is indicated. (**D**) Alignment of C-terminal peptide sequences of *Xbp1u* from fruit fly, zebrafish, mouse and human. The amino acids in the P-site position where the stalling occurs are indicated in bold. Conserved amino acids are highlighted in yellow, while the ones which are most likely critical for stalling are indicated with triangles. A pink triangle indicates a position where S -> A mutation has been demonstrated to increase the translational pausing [61].

### CHX causes initiation bias, while flash-freezing captures terminating ribosomes

The use of translational inhibitors has been shown to cause abnormalities in ribosome profiles. Specifically, in cycloheximide-treated samples, the slow diffusion of the drug allows translation to continue for a few codons before the CHX reaches 100% efficiency. This can be observed as an artifactual *‘*ramp*’* at the 5*’*end of the coding sequences [24, 26]. We observe the accumulation of initiating ribosomes in CHX-treated samples (Y1, Y2, Z3, Z5, Z7, M1, M3, H1, H3, H4), as well as those treated with emetine and rapamycin (F1, F2, F3). Interestingly, in samples where no drug was used or they have been flash-frozen before additional treatment with CHX (Y3, Z1, Z2, Z4, Z6, M2, H2), we observe minimal accumulation of ribosomes at the start codon, they however accumulate at the last sense codon of the CDS. This peak before stop codon likely comes from termination pausing, which allows time for the termination complex to assemble, release the peptide and dissociate. This termination peak is lost in CHX treated samples, due to the long time in the translational timescale required for CHX to reach equilibrium [24] which is likely too long to capture terminating ribosomes.

### Detection of robust and conserved stall sites to control for library bias

To obtain high-confidence stall sites (Figure 1A) we devised a method to detect stalls above the noise level and required these sites to be detected in at least two libraries from different organisms (see Methods). As most ribosome profiling experiments have used CHX to *‘*freeze*’* elongating ribosomes, this might skew the distribution of stall sites towards longer-lasting pauses. While flash-frozen samples may provide a more accurate picture of short-lived pauses, there is currently not enough data of this type to detect transient pausing above noise levels. We, therefore, limited our analysis to long-lasting pauses (1-2 seconds, given a mean decoding rate of 5.6 codons per second [29] and a slow inhibition by CHX allowing time for ribosomes to run-off for several codons [26]). Detection of such pauses should therefore be independent of the use of translation inhibitors and, given the support of multiple libraries, can be separated from artificial peaks produced by library-specific biases. Using this strategy we identified thousands of peaks in each of the libraries (see Supplementary Table 2), with only a small fraction of them replicated across experiments (yeast: 781; fruit fly: 1096; zebrafish: 167; mouse: 577; human: 674).

Stall sites that have biological significance are likely to be evolutionary conserved. We, therefore, compared the positions of peaks in homologous genes across all five analyzed organisms (see Materials and methods). This analysis revealed 3293 stall sites conserved in at least two organisms (Figure 1B, Supplementary Table 3). For human, we detected 2426 peaks in 1729 genes that are present in at least one other organism, and we will refer to these as *‘*conserved stall sites*’* (CSS). The highest degree of conservation is unsurprisingly seen over the shortest evolutionary distance, between mouse and human homologs. These sites account for ∼9% and ∼18% of total peaks for mouse and human, respectively. Surprisingly, we find some pause sites conserved even in yeast, which is evolutionarily distant from the other organisms. For the remaining organisms the number of stall sites that are conserved only accounts for 1-4% of all peaks (Supplementary Table 2). Since genes have a median length of around 500 codons and contain an average of ∼1.4 stall sites per gene, the probability of finding peaks in homologous genes at the same positions at random is very low. The high degree of conservation of stall sites therefore supports the idea that stalling is a non-random event and that the sites that we identify bear biological significance.

Stalling has been predicted from ribosome profiling previously [12]. While most of the studies focused on bacteria or yeast, to date only Ingolia et al. 2011 looked at genome-wide stalling in higher eukaryotes (mouse) [29]. Importantly, the pause sites in this and other studies are generally identified by the peak in read density alone and often in a single library. The seminal study of Ingolia et al. 2011 reported 1500 strong ribosomal pauses, evolutionarily conserved and estimated to last for several seconds. Since pauses of this duration should be captured independently of the method used for translation inhibition, we compared the peaks derived from this study (M2) that did not use translation inhibitors with CHX-treated libraries from the same study (M1) using our peak calling strategy. With our method we recovered all 1500 peaks reported previously together with 16474 of novel sites in M2. However, when comparing these with the CHX-treated library M1, the overlap is only 308 sites that are present in both libraries. To further investigate whether the subset of stall sites repeated in both libraries was in agreement with the sequence motif reported to cause stalling [29], we analyzed the peptide sequence around these stall sites. As our methodology for calling peaks is largely similar to Ingolia et al. 2011 (see Materials and methods), the peaks found in library M2 were consistent with the previous report with a strong enrichment of glutamate or aspartate in the A-site preceded by a proline or glycine and then another proline (Supplementary Figure 2A). However, identical analysis for pauses present in the overlap of both M1 and M2 libraries revealed that the P-site is most likely to contain an aspartate or glycine (Supplementary Figure 2B). The bias was similar for the A-site, though less pronounced. Both stall sites also revealed an influence from double prolines, which has previously been shown to cause stalling [3]. The low fraction of pause sites that is consistent across treatments and a changed sequence motif suggest that a significant number of sites reported previously are likely due to library bias and do not carry biological significance, though some could occur due to shorter transient pausing.

**Fig. 2:**
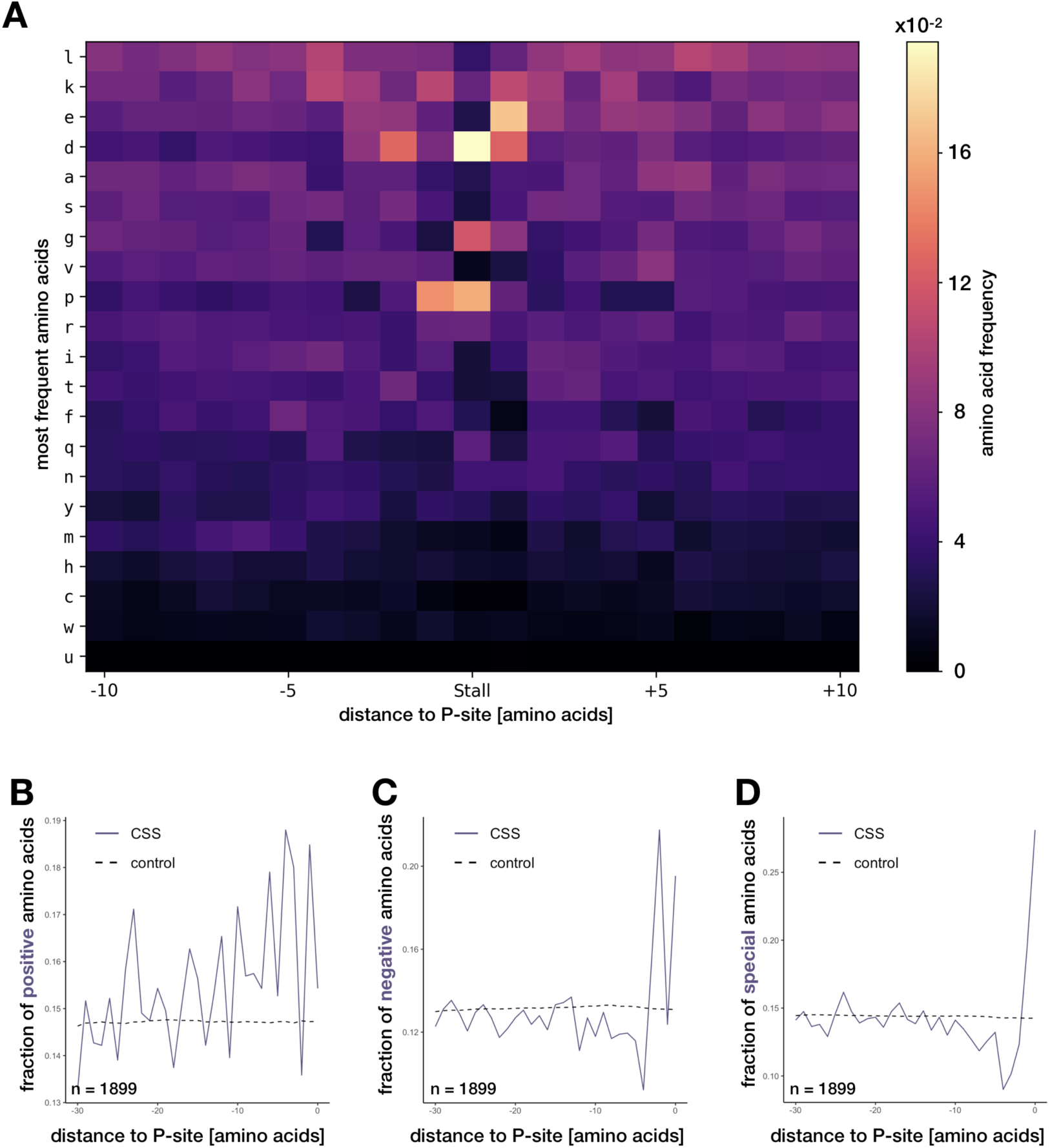
Mechanisms of conserved stalling. (**A**) Overrepresentation of amino acids around CSSs, sorted from most to least frequent, top to bottom. (**B**) Fraction of positively charged amino acids in the 30 amino acids upstream of the CSSs at P-site (0), spanning the exit tunnel versus background. Similarly, (**C**) fraction of negatively and (**D**) special (proline and glycine) amino acids.

A well-characterized example of stalling within a eukaryotic CDS is the transcription factor *XBP1*. During impairment of protein folding in the endoplasmic reticulum (ER), commonly known as ER stress, the nascent chain of *XBP1u* (the shorter, unspliced isoform of the transcript) localizes to the ER membrane and stalls. While stalled, the spliceosome on this membrane cuts out a fragment of yet untranslated mRNA, changing the open reading frame of the transcript and as a result produces an extended protein [61]. Mutational and evolutionary analysis of *XBP1u* peptide revealed peptide module at the carboxyl terminus required for pausing and splicing, of which 15 amino acids were conserved in human, mouse, chicken, frog and zebrafish and deemed necessary for stalling [61]. The exact position of stall site has been identified in mouse ribosome profiling data as a high peak over Asn256 codon in the A-site [29], corresponding to Met255 in the P-site. In our analysis we identified peaks close to the 3*’*-end of the *Xbp1u* mRNA in all of the libraries in the organisms where the two isoforms exist (fruit fly, zebrafish, mouse and human; Figure 1C and Supplementary Figure 3). The position of the peak is at the Met in P-site, with Asn in the A-site in zebrafish, mouse, and human. Interestingly, in fruit fly the identity of the P- and A-site codons is different, yet the peak occurs at the same position, as revealed by multiple sequence alignment (Figure 1D). In our analysis, the addition of fruit fly narrows down the number of conserved residues in the nascent peptide from 15 found before [61] to 5 amino acids, Pro in position −2 (relative to P-site at 0), Trp at −4, Glu at −7, Gly at −10 and Trp at −11. At position −5, the substitution of Ala (as in fruit fly) for Ser (as in the remaining organisms) has been shown to augment pausing [61]. Therefore, we conclude that these six residues are most likely critical for stalling on *Xbp1u*.

**Fig. 3:**
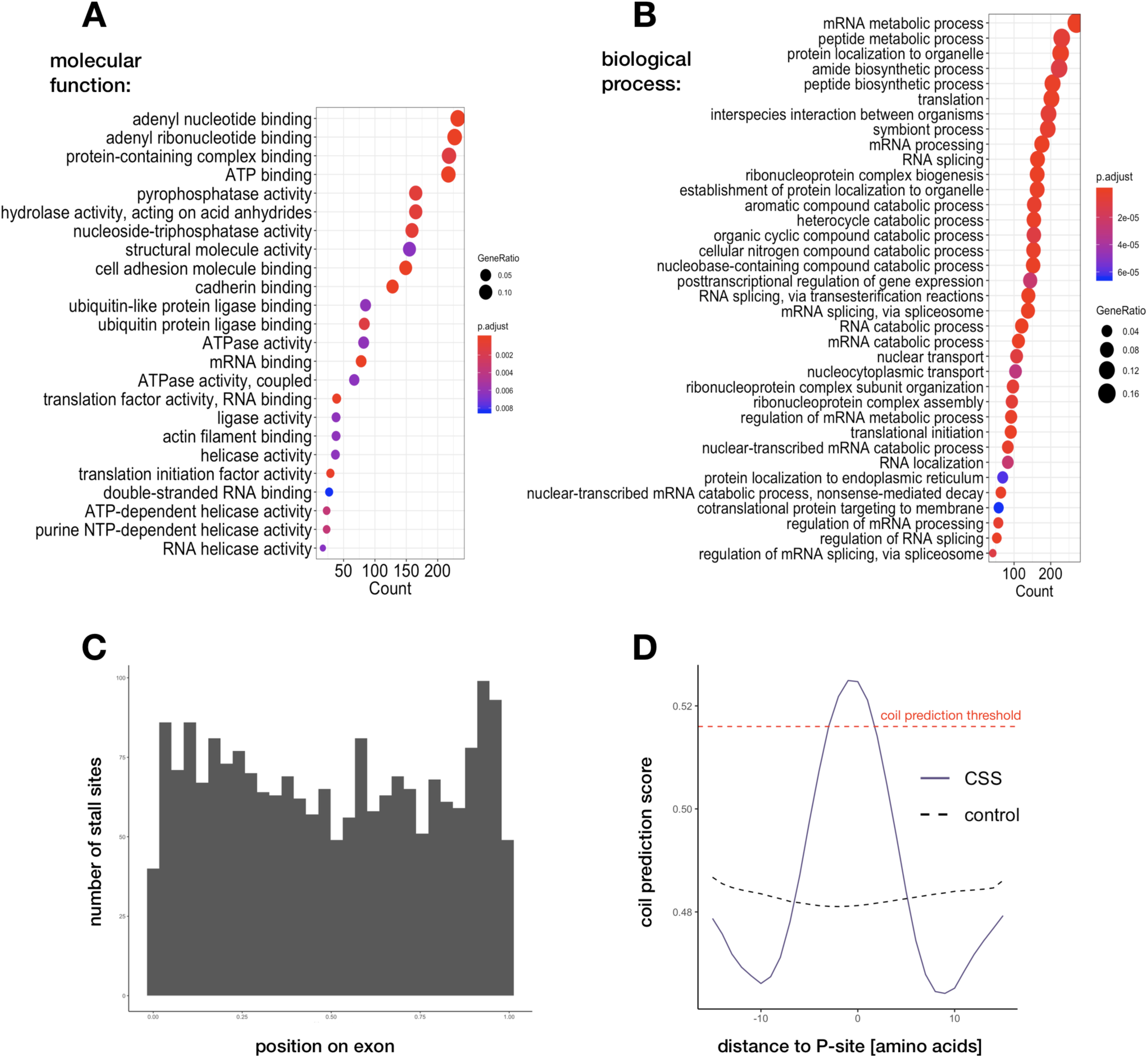
Functional characterization of CSS-containing genes. (**A**) and (**B**) Gene ontology enrichment analysis showing molecular functions and biological processes of conserved genes that are affected by stalling. (**C**) Position of CSSs on exons, with exon lengths normalized to 1. (**D**) Coil prediction score around CSSs, compared to control. Red dashed line marks the threshold typically used for coil prediction.

### Proline, glycine and negatively charged amino acids are conserved mechanisms of stalling

We investigated whether the organism-specific stall sites as well as CSSs are associated with factors that have been previously implicated in stalling by analyzing sequence and structure patterns around stall sites (see Methods). We corroborate the results that found proline as a major contributor to stalling [3]. Single proline amino acid in the P-site seems to indeed be one of the most influential individual contributors to conserved stalling, accounting for around 15% of CSSs (compared to around 6% of all codons in human coding for proline). The other significant contributors at P-site seem to be glycine (present in 12% CSSs, 7% in background) and aspartic acid (in 17% CSSs, 5% in background). Influential is also glutamic acid at A-site, found in 17% CSSs vs 7% in background, of which 10% do not overlap with Pro/Gly/Asp in P-site (Figure 2A).

We next asked whether similar biases could be observed in the nucleotide sequence by making a consensus logo of all CSSs (Supplementary Figure 4A). This revealed a bias towards a [CGA][CGA]N motif, which can be attributed to the codons for the enriched amino acids (proline: CCN, glycine: GGN, aspartic acid: GA[UC] at the P-site and glutamic acid: GA[AG] at the A-site). To investigate whether we could find any additional biases, we made individual logos for each of these amino acids (Supplementary Figure 4B). Together, 54% of all CSSs can be attributed to amino acid sequence, while the remaining 46% do not form any significant motif, neither in sequence logo nor by motif discovery (MEME) analysis [4]. Importantly, the sequence logos split by amino acids revealed nucleotide context in codons at positions −3 to +1. Such bias is not found when looking at other Pro, Gly, Asp, and Glu amino acids from CSS-containing genes that are not stall sites (not shown). To see if this sequence context can be explained by stretches of particular residues, we searched for amino acid 2-mers and 3-mers (Supplementary Figure 5). We do find significant stalling on proline, especially when present in doublets or triplets with another amino acid (XPP/PPX) [46]. These and other k-mers can explain up to 5% of CSSs, however, they occur at positions −1 to +1, which does not account for bias at position −3 to −2. An explanation for the latter could be that this sequence context is recognized by the ribosome, in a mechanism similar to recognition of Kozak sequence at the translation initiation sites [1], as the two contexts are of similar intensity.

A controversial question has been whether the charge of amino acids plays a role in stalling. While some studies have claimed that newly synthesized, positively charged amino acids contribute to stalling [14], others have disputed this [3] or only found a subtle effect and only in the absence of translation inhibitors [47]. Others again found negatively charged amino acids contributing to stalling under certain conditions [49]. Given the conservation of the translational machinery, it is likely that a charge-dependent mechanism will be conserved between multiple organisms. We therefore analyzed the 30 amino acids upstream of stall sites that would span the ribosome exit tunnel. Using random sites in the same gene as a control we found a small contribution from positively charged amino acids present immediately upstream of the ribosome active site, particularly at positions −1 (E-site), −3, and −4. The proportion of CSSs that have these sequences however is small (2.3% of stalling cases) (Figure 2B) and can be fully attributed to consecutive lysine codons, which are present in nearly 3% of sequences just upstream of CSSs, which are known to cause stalling [2, 35]. Consecutive lysine residues are indeed the most frequent 2-mers and 3-mers that influence stalling (see Supplementary figure 5A,B).

Interestingly, we find a much stronger contribution from negatively charged amino acids at position −2 (first amino acid in the exit tunnel), see Figure 2C. Most of them co-occur with the four amino acids found to be the strongest stalling contributors, especially Gly (34% of Gly-related CSSs; for others, the percentages are Asp: 22%, Pro: 20%, Glu: 14% and the rest: 15%). Altogether, in total, 12% of all CSSs are Gly/Pro/Asp/Glu-associated and have a negatively charged amino acid at position −2, additional 7% have the charge at −2, but do not have these amino acids. As the entrance to the exit tunnel is narrow [19], we hypothesize that upon encountering it, the negative charge of amino acids might repel the negative charge of the exit tunnel and slow down translation. Interestingly, the negatively charged amino acids at position −2 and positively charged at −1, −3, and −4 are never found together. Finally, as already observed, we see a large contribution from ?special? amino acids, proline and glycine, at the P-site, that add up to 27% CSSs (Figure 2D).

Overall, we can explain 63% of CSSs by sequence features: 44% by amino acids at P-site (15% Pro, 12% Gly and 17% Asp), 10% by glutamate at A-site, 2% by lysine stretches and additional 7% by negatively charged amino acids at the entrance to the exit tunnel.

mRNA structure has been shown to have an influence on translation slowdown [59, 58], and a recent study implicated its role in pausing of chloroplast ribosomes [23]. Gawroński et al. observed increased stability of mRNA secondary structure 31 nucleotides downstream of pausing site, for a sample of 78 stall sites. We investigated whether we could see any influence of structure in the CSSs that were not explained by sequence features. We analyzed *in silico* folded mRNAs, looking at the minimum free energy (MFE) around CSSs using the same method as previously. Our sample size was 627 CSSs, so much larger than in the chloroplast study. We did not find any decrease in MFE downstream of CSSs (Supplementary Figure 6). Therefore we conclude that mRNA secondary structure is unlikely to be a major cause of conserved stalling.

Synonymous single-nucleotide polymorphisms (SNPs) have been suggested to potentially induce stalling [57]. While a synonymous SNP does not alter the amino acid sequence, it can change the codon to a one that is rarely used. Such rare codons could provoke stalling [10, 48] due to lower concentrations of the cognate tRNAs. To investigate this possible association we searched *de novo* for SNPs as well as used the human SNP databases, containing over 36 million unique SNPs. We found no significantly different association of SNPs with stall sites as compared to random controls (see Methods), indicating that stall sites are not generally associated with a higher incidence of SNPs.

### Genes undergoing stalling are involved in RNA metabolism and co-translational protein folding

To investigate whether stalling plays a role in specific cellular processes we sought to determine whether the genes containing CSSs share common functions. Compared to a background of 4860 highly expressed genes, out of the 1729 CSS-containing genes, 1415 are associated with biological process ontology terms and 897 with molecular function terms. The genes are enriched in functions related to RNA metabolic processes, mRNA splicing and processing, but also protein targeting to the endoplasmic reticulum (as in the case of *XBP1*), translation regulation, and cotranslational protein targeting to the membrane. We also find terms related to nonsense-mediated decay which have been suggested as possible reasons for stalling [61, 22, 32, 13, 31] (see Figure 3A, 3B, Supplementary Table 2 and Supplementary Table 4). A similar analysis of 2064 well-expressed homologs (defined as having a median codon coverage higher than zero, see Methods) without CSSs against the same background returned no significant GO terms. This argues that stalling serves a specific function in the cell regulating a subset of genes involved in RNA metabolism and protein targeting.

Given the overrepresentation of genes involved in mRNA splicing in the conserved stall sites, we asked whether alternative splicing and stalling might regulate each other. We analyzed 200 bases of sequence flanking CSS to identify enrichment of splicing factor binding sites that potentially could be involved in regulating stalling if they remained bound to processed mRNA in the cytoplasm. As expected for exonic regions, we observed a general increase in GA rich sequences, but no specific splicing factor binding sites were identified (Supplementary Figure 7). When comparing the positioning of human CSSs relative to the closest splice site we found that stall sites tend to be positioned closer to the exon junctions, especially at the 3*’*end of exons (Figure 3C). This proximity to junctions could mean for instance that the stalls are there to allow time for unconventional cytoplasmic alternative splicing (UC-AS) [11], as in the case of *XBP1*, or indicate a possible coupling of conventional nuclear alternative splicing and NMD (AS-NMD). Such a mechanism has been proposed to act as post-transcriptional gene control, where alternative splicing creates a premature termination codon, targeting the resulting mRNA for NMD. Studies have indicated that many splicing regulatory proteins are themselves regulated by AS-NMD [43]. Another explanation could be that, as secondary structure elements tend to be contained within exons [40], stall sites at the ends of exons are there to allow the newly synthesized regions to order themselves, as suggested before [56, 34, 63]. To test this explanation, we predicted the level of disorder in the CSS-containing proteins using DisEMBL [39] (see Methods). This analysis revealed that CSSs tend to be located within coils, which are the linkers between alpha-helices and beta-strands (Figure 3D) as often as in 65% cases (with the average 55% for the random sites sampled from the same proteins). To see if similar dependence exists for tertiary structure elements, we analyzed distance to the closest upstream protein domain downloaded from the CATH database. However, we found no evidence that CSSs are more likely located downstream of higher-order domains than random (Supplementary Figure 8). Overall, a subset of CSSs might be involved in the co-translational folding of secondary structure protein domains.

### Little evidence for involvement of CSSs in membrane targeting and degradation

Stalling has been shown to facilitate co-translational protein targeting to the membrane [15, 44]. In a recent study, a significant pausing signal was observed downstream of the start of transmembrane domains (TMs) in chloroplasts. It occurred 52 amino acids downstream of the start of type II TMs and 34 amino acids for type I TMs [23]. The authors speculated that this would leave the time for the TMs to fold before translation would proceed. To investigate whether this type of stalling is a conserved mechanism we downloaded all 1512 TM type I and 464 type II proteins available in UniProt for human. Out of these, only 76 and 36 contained CSSs, respectively. The CSSs that were present in the TM proteins were distributed randomly over the body of the gene, and not overrepresented at any specific position downstream of the start of TMs. Therefore, we conclude that conserved pause sites are not associated with folding of transmembrane domains.

Some stall sites might be a consequence of aberrant translation. To investigate this hypothesis we sought to understand whether stalling could lead to abortion of translation. We therefore calculated the ratio of mean ribosome coverage upstream versus downstream of all CSSs in the four human libraries. We found that for the majority of stall sites the ratio is less than 2-fold, and only 22 have log2 ratio higher than 2 (see Supplementary figure 9). From manual inspection of these we found 13 stall sites in 11 genes that might lead to programmed abortion of translation, possibly by NMD or NGD (Supplementary Table 5), as these were located over out-of-frame stop codons or just before alternative exons.

Similarly, knowing that canonical termination of translation produces longer footprints due to conformational change of the ribosome [29], we investigated whether we could find longer footprints at CSSs if these led to termination or different fragment length distribution whatsoever. Indeed, we observe a shift in average footprint length around stop codons in the four human libraries (Supplementary figure 10A). However, no such shift is observed around CSSs present in these libraries (Supplementary figure 10B). Overall, we conclude that CSSs do not tend to lead to abortion of translation or conformational change of the ribosome.

## Discussion

The aim of this study was to find and characterize conserved functional stall sites. Using a representative set of libraries with good coverage for five model organisms, we tailored processing to each dataset separately to allow for robust comparison. We identified thousands of library-specific peaks, of which 3293 stall sites were conserved in at least two organisms.

While many mechanisms have been suggested to induce stalling we found that 63% of conserved stalling can be explained by proline, glycine, and negatively charged amino acids given the proper context. While we cannot exclude that RNA structure contributes to stalling it does not seem to play a significant part at our CSSs.

Interestingly, many of the CSSs identified in this study are present in genes coding for RNA processing factors, notably splicing factors. The mechanistic significance of this at a global level is yet unclear and any possible connections between regulation of unconventional cytoplasmic splicing such as in the case of *XBP1* and stalling are not immediately obvious. Another interesting category includes proteins with functions such as co-translational folding or membrane targeting, which are thought to be regulated by stalling. We do find that CSSs tend to be located in between ordered protein domains, suggesting a role in co-translational folding. However, as such proteins are involved in these processes themselves, this might imply a possible self-regulation mechanism, where stalling regulates the synthesis of such proteins, but in turn, the synthesized proteins regulate the stalling during translation. This is an attractive hypothesis for which our study provides candidates for further experimental testing.

Given the high variability in terms of cell lines and tissues of the data analyzed here, it is likely that the conserved stalling landscape is larger than the sites identified in this study and includes stall sites that are specific to certain conditions. Also, the vast majority of ribosome profiling experiments perform size-selection, keeping only the footprint sizes of 28-29 nucleotides, typical of the non-rotated, elongating ribosomes, but it has been shown that lengths one might expect at stall sites include those from closely stacked di-ribosomes protecting around 80 nucleotides [25] or short footprints 20-22 nucleotides long representing the rotated form of the ribosome [37]. Therefore keeping longer fragments might be beneficial for analysis of stalling.

In conclusion, this study presents a rich resource on global, conserved ribosome stall sites, indicating possible causes and implications. The methods and data presented here lay the foundation for further research involving in-depth molecular biology to characterize the functional relevance of the identified conserved stall sites.

## Materials and methods

### Ribosome profiling data

The raw sequencing data were downloaded from the Sequence Read Archive (SRA). The normal condition libraries include yeast: SRR387905 [27], SRR1520317 cycloheximide (CHX)-treated and SRR1520325 (no drug) [24]; fruit fly S2 cells: SRR942880 [21], SRR6930625 [42] and SRR3031135; zebrafish embryos at different stages of development: 2 hours post fertilization (hpf) - SRX399824, SRX399826, SRX399828 [6], SRR5893147, SRR5893148 [7] and SRR1039873 [55], 4 hpf - SRR836195 [16] and SRR1039876 [55], 6 hpf - SRR836196 [16] and SRR1039879 [55]; mouse: embryonic stem cells - SRR315601, SRR315602 (CHX-treated), SRR315616, SRR315617, SRR315618, SRR315619 (no drug) [29] and 3T3 fibroblasts - SRR1039863 [55]; human: fibroblasts - SRR609197 (CHX-treated) and SRR592961 (no drug) [53], mitotic HeLa cells - SRR970587 [54] and HEK293 cells - SRR1039861 [55]. Overview of data libraries is shown in Supplementary Table 1.

### Ribosome profiling data processing

Each of the raw ribosome profiling sequencing libraries was processed in the following way: (1) trimming adapters specific for each experiment and low quality bases with *cutadapt*, keeping reads of minimum length of 20 nucleotides; (2) removing reads mapping to ribosomal RNAs from *SILVA* rRNA database and organism-specific non coding RNAs from *Ensembl* using *Bowtie2*; (3) aligning the remaining reads to organism-specific reference transcriptome with *TopHat2*, allowing up to 2 mismatches. The *Ensembl* genome versions used were R64-1-1 for yeast, BDGP6 for fruit fly, GRCz10 for zebrafish, GRCm38 for mouse and GRCh38 for human; (4) selecting the periodic footprint lengths and assigning them to P-site nucleotides with Shoelaces [8]. The selected lengths and offsets are shown in Supplementary Table 1. The ribosome meta-profiles (Supplementary Figure 1) were plotted by taking the last 30 nucleotides of 5*’*UTR, first and last 60 nucleotides of CDS and 30 first nucleotides of 3*’*UTR (from the genes that contain features of minimum those lengths) and superimposing them, taking into account the length of the footprint.

### Stall site calling

Ribosome footprint profiles for each transcript were constructed by quantifying the number of footprints assigned to each nucleotide position. For organisms with multiple transcripts per gene (all but yeast), only the longest transcripts were used. For prediction of stall sites, a further subset of well-expressed transcripts, defined as having a median codon coverage higher than zero, were selected. As there are usually high peaks over start and stop codons due to prolonged time of initiation and elongation, the first and the two last codons of each CDS (start, stop and a codon before stop codon) were excluded from further analysis to avoid skewing the footprint distribution over transcripts. The codon coverage per transcript was then transformed into z-scores, and the stall sites were identified as codons with coverage higher than a certain threshold (experimental cut-off of 5.0 was chosen). The peaks within first 5 codons of CDSs were excluded, to avoid these caused by accumulation of ribosomes at the beginning of CDSs in some libraries. To further increase the confidence that defined peaks are indeed stall sites and not experimental or sequencing biases, the peak had to occur in at least two different organisms (see below) to be considered a stall site. The number of peaks in each library and overlap between libraries is shown in Supplementary Table 2.

### Conservation analysis

Sets of homologous genes for each organism were retrieved from *Ensembl* using *Biomart* querying. The transcript sequences for each set of homologs were aligned together using *Clustal Omega*. The positions of stall sites for each organism were then cross-checked among homologous transcripts to account for insertions and deletions. If the stall site occurs at the same or adjacent codon in homologs in given organisms, it is considered to be conserved in these. The list of conserved stall sites is presented in the Supplementary Table 3.

Note that *Xbp1* is not listed in the results table, as it has been generated based on alignments of longest transcript per gene from *Ensembl* annotations. Manual inspection revealed that the correct transcripts of *Xbp1s/u* for mouse and humans are missing in *Ensembl*, which shifted the alignments. Therefore these had to be downloaded from RefSeq, and the shorter *Xbp1u* transcripts have been aligned and analyzed as others.

### SNP analysis

For frequent to rare substitution analysis, *de novo* SNPs in the H2 library (which has the highest coverage among the human libraries) were called with bcftools [38] with default settings, predicting 14 SNPs with low coverage. None of them were associated with CSSs.

Human CSSs were then checked for association with all known single nucleotide polymorphisms (SNPs) from dbSNP database [33], clinically associated SNPs from ClinVar database [36], and also the 14 novel SNPs from the H2 library. For each stall site, a random codon on the same gene has been used as control. We estimated that the chance of finding a SNP at a specific position in the CDS regions (see Stall site calling) is 0.009. Therefore, the expected count of SNPs overlapping with control stall sites or conserved stall sites is 65 (dbSNP, ClinVar). In the H2 library we find 55 (dbSNP) and 54 (ClinVar) SNPs that overlap with control stall sites, and 54 (dbSNP) and 53 (ClinVar) that overlap with conserved stall sites. Restriction to only non-synonymous SNPs results in 27 (dbSNP) and 31 (ClinVar) over control stall sites, while 35 (dbSNP) and 33 (ClinVar) over conserved stall sites.

### Sequence analysis

For amino acid heatmaps, we calculated the frequency of occurrence of specific amino acids and their 2-mers and 3-mers in the positions 10 amino acids upstream and downstream of CSSs.

For nascent peptide analysis, the 30 amino acids upstream of CSSs that would span the ribosome exit tunnel, were summed up depending on properties (positively charged, negatively charged, special). For control, we sampled 10000 times a random position from each of the transcripts with CSSs and calculated the average.

Sequence logos were created with *WebLogo3* [18] for all CSSs, as well as split by the most common amino acids. Additionally, we performed motif discovery with MEME suite [4], but found no reliable result.

### Structure analysis

In silico mRNA secondary structures of transcripts were predicted by calculating a minimum free energy (MFE) in a 51-nucleotide sliding window over CDSs using RNAfold program [41], as done previously [50, 23]. For structure analysis we used only stall sites positioned more than 30 nucleotides downstream from the start codon and 60 nucleotides upstream from the stop codon to avoid the decreased structure at the beginning and end of CDSs impacting the results of the analysis. This left 627 CSSs which were not explained by sequence features. As control, for each stall site we picked at random a position in the given transcript that neither overlaps with the region around the stall site (−30/+60 bases) nor the regions around start/stop codon as mentioned above. This was repeated 10000 times with different random positions and averaged. We over-imposed averaged regions around CSSs and average control in a meta plot.

### Gene Ontology enrichment

The genes with conserved stall sites present in human and at least one other organism were subject to Gene Ontology enrichment analysis. The enrichment was performed with cluster-ProfileR package [62] against a background of well-expressed homologs, as the set of stall sites was biased towards well-expressed homologs as well. As a control, we used well-expressed homologs without peaks (maximum z-score < 5 or 4, see Supplementary Table 2).

### Exon and protein domain boundary analysis

For splicing factors binding sites, we analyzed 200 nucleotides flanking CSSs for 6-mer content. Given the annotated splice junctions of the transcripts, we calculated the distance from CSSs in nucleotides to the upstream and downstream exon boundaries, and relative position of CSSs within the exons, excluding the first 45 and last 3 nucleotides. For protein domain boundary analysis, we downloaded protein domain annotations from CATH database [20]. We analyzed 1317 CSSs on 1100 genes that had an annotated domain, calculating the minimum distance to the upstream C-terminal protein domain boundary. For control, we used the random positions on the same genes.

To predict disorder in CSS-containing proteins, we used DisEMBL [39]. We extracted disorder scores for prediction of loops/coils around CSSs and averaged in a meta-plot. For control, we performed random selection of amino acids on the same proteins, excluding first 15, last 2 and those corresponding to CSSs, repeating 100 times for every gene. Additionally, we calculated the percentage of CSSs found in coils, as defined by default threshold of 0.516 versus average percentage of random sites.

### Transmembrane domain analysis

We downloaded all 1512 TM type I and 464 type II proteins available in UniProt for human (annotation SL-9905 and SL-9906). Out of these, we selected those containing CSSs, resulting in 76 and 36 for type I and type II, respectively. We looked at the distribution of these in the genes, and found no overrepresentation at any certain position downstream of the signal peptide.

### Premature termination analysis

For the human libraries, we analyzed transcripts that contained CSSs and had at least 15 codons before and after the CSS contained within the body of the gene (excluding first 15 and last 2 codons). To check for reduction of signal 3*’* of the pause site, we calculated log2 ratio of the mean ribosome coverage in the upstream region to the mean coverage in the downstream region from the CSS. For fragment length distribution, for libraries H1-H4 we calculated background distributions of all footprints. We extracted coverage 15 nt upstream and 15 nt downstream around all stop codons (for the transcripts that contain 3*’*UTRs) and created metaplots of average footprint lengths at these positions. Similarly, for each of the libraries, we extracted coverage around CSSs present in a given library and created metaplots as in the case of stop codons.

## References

[1] Acevedo, J. M., Hoermann, B., Schlimbach, T., and Teleman, A. A. Changes in global translation elongation or initiation rates shape the proteome via the Kozak sequence. Scientific Reports 8, 1 (Mar. 2018), 337.

[2] Arthur, L. L., Pavlovic-Djuranovic, S., Koutmou, K. S., Green, R., Szczesny, P., and Djuranovic, S. Translational control by lysine-encoding A-rich sequences. Science Advances 1, 6 (July 2015), e1500154.

[3] Artieri, C. G., and Fraser, H. B. Accounting for biases in riboprofiling data indicates a major role for proline in stalling translation. Genome Research 24, 12 (Oct. 2014), gr.175893.114–2021.

[4] Bailey, T. L., Boden, M., Buske, F. A., Frith, M., Grant, C. E., Clementi, L., Ren, J., Li, W. W., and Noble, W. S. MEME SUITE: tools for motif discovery and searching. Nucleic Acids Research 37, Web Server (June 2009), W202–W208.

[5] Bartholomäus, A., Del Campo, C., and Ignatova, Z. Mapping the non-standardized biases of ribosome profiling. Biological Chemistry 397, 1 (2016), 23–35.

[6] Bazzini, A. A., Johnstone, T. G., Christiano, R., Mackowiak, S. D., Obermayer, B., Fleming, E. S., Vejnar, C. E., Lee, M. T., Rajewsky, N., Walther, T. C., and Giraldez, A. J. Identification of small ORFs in vertebrates using ribosome footprinting and evolutionary conservation. The EMBO Journal 33, 9 (May 2014), 981–993.

[7] Beaudoin, J.-D., Novoa, E. M., Vejnar, C. E., Yartseva, V., Takacs, C. M., Kellis, M., and Giraldez, A. J. Analyses of mRNA structure dynamics identify embryonic gene regulatory programs. Nature Structural & Molecular Biology 25, 8 (Aug. 2018), 677–686.

[8] Birkeland, Å., Chyzynska, K., and Valen, E. Shoelaces: an interactive tool for ribosome profiling processing and visualization. BMC Genomics 19, 1 (Dec. 2018), 543.

[9] Brandman, O., and Hegde, R. S. Ribosome-associated protein quality control. Nature Structural ‘Molecular Biology 23, 1 (Jan. 2016), 7–15.

[10] Brule, C. E., and Grayhack, E. J. Synonymous Codons: Choose Wisely for Expression. Trends in Genetics 33, 4 (Apr. 2017), 283–297.

[11] Buckley, P. T., Khaladkar, M., Kim, J., and Eberwine, J. Cytoplasmic intron retention, function, splicing, and the sentinel RNA hypothesis. Wiley interdisciplinary reviews. RNA 5, 2 (Mar. 2014), 223–230.

[12] Buskirk, A. R., and Green, R. Ribosome pausing, arrest and rescue in bacteria and eukaryotes. Philos Trans R Soc Lond B Biol Sci 372, 1716 (Mar 2017).

[13] Celik, A., Kervestin, S., and Jacobson, A. NMD: At the crossroads between translation termination and ribosome recycling. Biochimie 114 (July 2015), 2–9.

[14] Charneski, C. A., and Hurst, L. D. Positively Charged Residues Are the Major Determinants of Ribosomal Velocity. PLoS Biology 11, 3 (Mar. 2013), e1001508.

[15] Chartron, J. W., Hunt, K. C. L., and Frydman, J. Cotranslational signal-independent SRP preloading during membrane targeting. Nature 536, 7615 (Aug. 2016), 224–228.

[16] Chew, G. L., Pauli, A., Rinn, J. L., Regev, A., Schier, A. F., and Valen, E. Ribosome profiling reveals resemblance between long non-coding RNAs and 5’ leaders of coding RNAs. Development 140, 13 (July 2013), 2828–2834.

[17] Collart, M. A., and Weiss, B. Ribosome pausing, a dangerous necessity for cotranslational events. Nucleic Acids Res 48, 3 (Feb 2020), 1043–1055.

[18] Crooks, G. E., Hon, G., Chandonia, J.-M., and Brenner, S. E. WebLogo: a sequence logo generator. Genome Research 14, 6 (June 2004), 1188–1190.

[19] Dao Duc, K., Batra, S. S., Bhattacharya, N., Cate, J. H. D., and Song, Y. S. Differences in the path to exit the ribosome across the three domains of life. Nucleic Acids Research 47, 8 (Feb. 2019), 4198–4210.

[20] Dawson, N. L., Lewis, T. E., Das, S., Lees, J. G., Lee, D., Ashford, P., Orengo, C. A., and Sillitoe, I. CATH: an expanded resource to predict protein function through structure and sequence. Nucleic Acids Research 45, D1 (Jan. 2017), D289–D295.

[21] Dunn, J. G., Foo, C. K., Belletier, N. G., Gavis, E. R., Weissman, J. S., and Sonenberg, N. Ribosome profiling reveals pervasive and regulated stop codon readthrough in Drosophila melanogaster. eLife 2 (Dec. 2013), e01179.

[22] Fluman, N., Navon, S., Bibi, E., and Pilpel, Y. mRNA-programmed translation pauses in the targeting of E. coli membrane proteins. eLife 3 (Aug. 2014), 693.

[23] Gawroński, P., Jensen, P. E., Karpiński, S., Leister, D., and Scharff, L. B. Pausing of Chloroplast Ribosomes Is Induced by Multiple Features and Is Linked to the Assembly of Photosynthetic Complexes. Plant Physiology 176, 3 (Mar. 2018), 2557–2569.

[24] Gerashchenko, M. V., and Gladyshev, V. N. Translation inhibitors cause abnormalities in ribosome profiling experiments. Nucleic Acids Research 42, 17 (July 2014), gku671–e134.

[25] Guydosh, N. R., and Green, R. Dom34 Rescues Ribosomes in 3’ Untranslated Regions. Cell 156, 5 (Feb. 2014), 950–962.

[26] Hussmann, J. A., Patchett, S., Johnson, A., Sawyer, S., and Press, W. H. Understanding Biases in Ribosome Profiling Experiments Reveals Signatures of Translation Dynamics in Yeast. PLoS Genetics 11, 12 (Dec. 2015), e1005732.

[27] Ingolia, N., Brar, G. A., Rouskin, S., Mcgeachy, A. M., and Weissman, J. S. The ribosome profiling strategy for monitoring translation in vivo by deep sequencing of ribosome-protected mRNA fragments. Nature Protocols 7, 8 (July 2012), 1534–1550.

[28] Ingolia, N., Ghaemmaghami, S., Newman, J. R. S., and Weissman, J. S. Genome-wide analysis in vivo of translation with nucleotide resolution using ribosome profiling. Science 324, 5924 (Apr. 2009), 218–223.

[29] Ingolia, N., Lareau, L. F., and Weissman, J. S. Ribosome Profiling of Mouse Embryonic Stem Cells Reveals the Complexity and Dynamics of Mammalian Proteomes. Cell 147, 4 (Nov. 2011), 1–14.

[30] Ito, K., and Chiba, S. Arrest Peptides: *Cis*-Acting Modulators of Translation. Annual Review of Biochemistry 82, 1 (June 2013), 171–202.

[31] Joazeiro, C. A. P. Ribosomal Stalling During Translation: Providing Substrates for Ribosome-Associated Protein Quality Control. doi.org 33, 1 (Oct. 2017), 343–368.

[32] Kim, J., Klein, P. G., and Mullet, J. E. Ribosomes pause at specific sites during synthesis of membrane-bound chloroplast reaction center protein D1. Journal of Biological Chemistry 266, 23 (Aug. 1991), 14931–14938.

[33] Kitts, A., and Sherry, S. The Single Nucleotide Polymorphism Database (dbSNP) of Nucleotide Sequence Variation. In The NCBI Handbook [Internet]. National Center for Biotechnology Information (US), Feb. 2011.

[34] Komar, A. A. A pause for thought along the co-translational folding pathway. Trends in Biochemical Sciences 34, 1 (Jan. 2009), 16–24.

[35] Koutmou, K. S., Schuller, A. P., Brunelle, J. L., Radhakrishnan, A., Djuranovic, S., and Green, R. Ribosomes slide on lysine-encoding homopolymeric A stretches. eLife 4 (Feb. 2015), 1–51.

[36] Landrum, M. J., Lee, J. M., Riley, G. R., Jang, W., Rubinstein, W. S., Church, D. M., and Maglott, D. R. ClinVar: public archive of relationships among sequence variation and human phenotype. Nucleic Acids Research 42, Database issue (Jan. 2014), D980–D985.

[37] Lareau, L. F., Hite, D. H., Hogan, G. J., and Brown, P. O. Distinct stages of the translation elongation cycle revealed by sequencing ribosome-protected mRNA fragments. eLife 3 (2014), e01257.

[38] Li, H. A statistical framework for SNP calling, mutation discovery, association mapping and population genetical parameter estimation from sequencing data. Bioinformatics 27, 21 (Nov. 2011), 2987–2993.

[39] Linding, R., Jensen, L. J., Diella, F., Bork, P., Gibson, T. J., and Russell, R. B. Protein Disorder Prediction: Implications for Structural Proteomics. Structure 11, 11 (Nov. 2003), 1453–1459.

[40] Liu, M., and Grigoriev, A. Protein domains correlate strongly with exons in multiple eukaryotic genomes – evidence of exon shuffling? Trends in Genetics 20, 9 (Sept. 2004), 399–403.

[41] Lorenz, R., Bernhart, S. H., Siederdissen, C. H. Z., Tafer, H., Flamm, C., Stadler, P. F., and Hofacker, I. L. ViennaRNA Package 2.0. Algorithms for Molecular Biology 6, 1 (Dec. 2011), 26.

[42] Luo, S., He, F., Luo, J., Dou, S., Wang, Y., Guo, A., and Lu, J. Drosophila tsRNAs preferentially suppress general translation machinery via antisense pairing and participate in cellular starvation response. Nucleic Acids Research 46, 10 (June 2018), 5250–5268.

[43] Mcglincy, N. J., and Smith, C. W. J. Alternative splicing resulting in nonsense-mediated mRNA decay: what is the meaning of nonsense? Trends in Biochemical Sciences 33, 8 (Aug. 2008), 385–393.

[44] Pechmann, S., Chartron, J. W., and Frydman, J. Local slowdown of translation by nonoptimal codons promotes nascent-chain recognition by SRP in vivo. Nature Structural and Molecular Biology (Nov. 2014).

[45] Pechmann, S., and Frydman, J. Evolutionary conservation of codon optimality reveals hidden signatures of cotranslational folding. Nature Structural & Molecular Biology 20, 2 (Feb. 2013), 237–243.

[46] Peil, L., Starosta, A. L., Lassak, J., Atkinson, G. C., Virumäe, K., Spitzer, M., Tenson, T., Jung, K., Remme, J., and Wilson, D. N. Distinct XPPX sequence motifs induce ribosome stalling, which is rescued by the translation elongation factor EF-P. Proceedings of the National Academy of Sciences 110, 38 (Sept. 2013), 15265–15270.

[47] Requião, R. D., De Souza, H. J. A., Rossetto, S., Domitrovic, T., and Palhano, F. L. Increased ribosome density associated to positively charged residues is evident in ribosome profiling experiments performed in the absence of translation inhibitors. RNA Biology 13, 6 (May 2016), 561–568.

[48] Rodnina, M. V. The ribosome in action: Tuning of translational efficiency and protein folding. Protein Science 25, 8 (June 2016), 1390–1406.

[49] Sabi, R., and Tuller, T. A comparative genomics study on the effect of individual amino acids on ribosome stalling. BMC Genomics 16, Suppl 10 (Oct. 2015), S5.

[50] Scharff, L. B., Childs, L., Walther, D., and Bock, R. Local Absence of Secondary Structure Permits Translation of mRNAs that Lack Ribosome-Binding Sites. PLoS Genetics 7, 6 (June 2011), e1002155.

[51] Shoemaker, C. J., and Green, R. Translation drives mRNA quality control. Nature Structural & Molecular Biology 19, 6 (June 2012), 594–601.

[52] Stein, K. C., and Frydman, J. The stop-and-go traffic regulating protein biogenesis: How translation kinetics controls proteostasis. Journal of Biological Chemistry 294, 6 (Feb. 2019), 2076–2084.

[53] Stern-Ginossar, N., Weisburd, B., Michalski, A., Le, V. T. K., Hein, M. Y., Huang, S.-X., Ma, M., Shen, B., Qian, S.-B., Hengel, H., Mann, M., Ingolia, N., and Weissman, J. S. Decoding Human Cytomegalovirus. Science 338, 6110 (Nov. 2012), 1088–1093.

[54] Stumpf, C. R., Moreno, M. V., Olshen, A. B., Taylor, B. S., and Ruggero, D. The translational landscape of the mammalian cell cycle. Mol Cell 52, 4 (Nov 2013), 574–582.

[55] Subtelny, A. O., Eichhorn, S. W., Chen, G. R., Sive, H., and Bartel, D. P. Poly(a)-tail profiling reveals an embryonic switch in translational control. Nature 508, 7494 (Apr 2014), 66–71.

[56] Thanaraj, T. A., and Argos, P. Ribosome-mediated translational pause and protein domain organization. Protein Science 5, 8 (Aug. 1996), 1594–1612.

[57] Tsai, C.-J., Sauna, Z. E., Kimchi-Sarfaty, C., Ambudkar, S. V., Gottesman, M. M., and Nussinov, R. Synonymous mutations and ribosome stalling can lead to altered folding pathways and distinct minima. J Mol Biol 383, 2 (Nov 2008), 281–291.

[58] Tuller, T., Veksler-Lublinsky, I., Gazit, N., Kupiec, M., Ruppin, E., and Zivukelson, M. Composite effects of gene determinants on the translation speed and density of ribosomes. Genome Biology 12, 11 (Nov. 2011), R110.

[59] Wen, J.-D., Lancaster, L., Hodges, C., Zeri, A.-C., Yoshimura, S. H., Noller, H. F., Bustamante, C., and Tinoco, I. Following translation by single ribosomes one codon at a time. Nature 452, 7187 (Mar. 2008), 598–603.

[60] Xie, P. Dwell-Time Distribution, Long Pausing and Arrest of Single-Ribosome Translation through the mRNA Duplex. International Journal of Molecular Sciences 16, 10 (Oct. 2015), 23723–23744.

[61] Yanagitani, K., Kimata, Y., Kadokura, H., and Kohno, K. Translational pausing ensures membrane targeting and cytoplasmic splicing of *XBP1u* mRNA. Science 331, 6017 (Feb. 2011), 586–589.

[62] Yu, G., Wang, L.-G., Han, Y., and He, Q.-Y. clusterProfiler: an R Package for Comparing Biological Themes Among Gene Clusters. OMICS: A Journal of Integrative Biology 16, 5 (May 2012), 284–287.

[63] Zhang, G., Hubalewska, M., and Ignatova, Z. Transient ribosomal attenuation coordinates protein synthesis and co-translational folding. Nature Structural & Molecular Biology 16, 3 (Mar. 2009), 274–280.

