## Supplementary Figures for "Deep conservation of ribosome stall sites across RNA processing genes"

The supplementary tables 2-5 are available in the XLSX file.

**Supplementary Table 1**: Data overview.

**Supplementary Table 2**: Peak statistics.

**Supplementary Table 3**: Conserved stall sites.

**Supplementary Table 4**: Gene Ontology biological processes and associated genes with CSSs.

**Supplementary Table 5**: Genes and stall sites that might lead to translation degradation.

**Supplementary Table 1**: Data overview. (*) Treatment with cycloheximide was performed *after* flash-freezing the embryos.

| Library name | Organism | Strain /cell line / stage | SRA accession | Translation inhibitor | Footprint lengths and offsets | | | | | | | | | | | | | | | | | | | | | Reference |
|---|---|---|---|---|---|---|---|---|---|---|---|---|---|---|---|---|---|---|---|---|---|---|---|---|---|---|
| | | | | | 20 | 21 | 22 | 23 | 24 | 25 | 26 | 27 | 28 | 29 | 30 | 31 | 32 | 33 | 34 | 35 | 36 | 37 | 38 | 39 | 40 | |
| **Y1** | **yeast** | **gb15** | SRR387905 | CHX | | 12 | 12 | 12 | 12 | 12 | 12 | 12 | 12 | 12 | 12 | 12 | 12 | 12 | 12 | | | | | | | Brar *et al.* 2012 |
| **Y2** | | **BY4741** | SRR1520317 | CHX | | | | | 8 | 9 | 10 | 11 | 12 | 13 | 14 | | | | | | | | | | | Gerashchenko *et al.* 2014 |
| **Y3** | | | SRR1520325 | no drug | | | | | | 9 | 10 | 11 | 12 | 13 | 14 | 14 | 14 | | | | | | | | | |
| **F1** | **fruit fly** | **S2 cells** | SRR942879 | emetine | | | | | | | 12 | 12 | 12 | 12 | 12 | 13 | 14 | 14 | 14 | 14 | 14 | 14 | 14 | 14 | 15 | Dunn *et al.* 2013 |
| **F2** | | | SRR6930625 | rapamycin | | | 8 | 8 | 8 | 11 | 11 | 12 | 12 | 13 | 13 | 13 | 14 | 14 | 14 | | | | | | | Luo *et al.* 2018 |
| **F3** | | | SRR3031135 | emetine | | 5 | 5 | 5 | 6 | 7 | 9 | 9 | 10 | 11 | 12 | 13 | 13 | 13 | 13 | 13 | | | | | | |
| **Z1** | **zebrafish** | **embryo, 2hpf** | SRX399824, SRX399826, SRX399828 | CHX* | 3 | 4 | 5 | 6 | 7 | 8 | 9 | 10 | 12 | 12 | 12 | 13 | | | | | | | | | | Bazzini *et al.* 2014 |
| **Z2** | | | SRR5893147, SRR5893148 | CHX* | | | | | | 9 | 10 | 11 | 12 | 12 | 13 | 14 | | | | | | | | | | Beaudoin *et al.* 2018 |
| **Z3** | | | SRR1039873 | CHX | | | | | | | 12 | 12 | | | | | 12 | | | | | | | | | Subtelny *et al.* 2014 |
| **Z4** | | **Embryo, 4hpf** | SRR836195 | CHX* | | | | | | 8 | 9 | 11 | 11 | 12 | 12 | 13 | | | | | | | | | | Chew *et al.* 2013 |
| **Z5** | | | SRR1039876 | CHX | | | | | | | | | | | 12 | | | | | | | | | | | Subtelny *et al.* 2014 |
| **Z6** | | **embryo, 6hpf** | SRR836196 | CHX* | | | | | | 8 | 8 | 11 | 11 | 12 | 12 | 13 | 13 | | | | | | | | | Chew *et al.* 2013 |
| **Z7** | | | SRR1039879 | CHX | | | | | | | | | | | 12 | | | | | | | | | | | Subtelny *et al.* 2014 |
| **M1** | **mouse** | **ESC** | SRR315601, SRR315602 | CHX | | 3 | 3 | | | | | 12 | 12 | 12 | 12 | 13 | 14 | 14 | 14 | 14 | | | | | | Ingolia *et al.* 2011 |
| **M2** | | | SRR315616, SRR315617, SRR315618, SRR315619 | no drug | | | | | | 7 | 8 | 9 | 10 | 11 | 12 | 13 | 14 | 14 | 14 | 14 | | | | | 14 | |
| **M3** | | **3T3** | SRR1039863 | CHX | | | | | 11 | 11 | 11 | 12 | 13 | 13 | 13 | 13 | 13 | 13 | 13 | 13 | | | | | 14 | Subtelny *et al.* 2014 |
| **H1** | **human** | **fibroblasts** | SRR609197 | CHX | | | 5 | 5 | 5 | 8 | 8 | 8 | 11 | 12 | 12 | 13 | 13 | 14 | 14 | 14 | 14 | | | | 14 | Stern *et al.* 2012 |
| **H2** | | | SRR592961 | no drug | | 3 | 4 | 5 | 6 | 7 | 8 | 9 | 10 | 11 | 12 | 13 | 14 | 14 | 14 | 14 | | | | | 14 | |
| **H3** | | **HeLa** | SRR970587 | CHX | 3 | 4 | 5 | 6 | 6 | 7 | 8 | 9 | 12 | 12 | 12 | 13 | 13 | 14 | 14 | 14 | | | | | | Stumpf *et al.* 2013 |
| **H4** | | **HEK293** | SRR1039861 | CHX | | | | | | 13 | 13 | 13 | 13 | 13 | 13 | 13 | 13 | 13 | 13 | 13 | | | | | | Subtelny *et al.* 2014 |

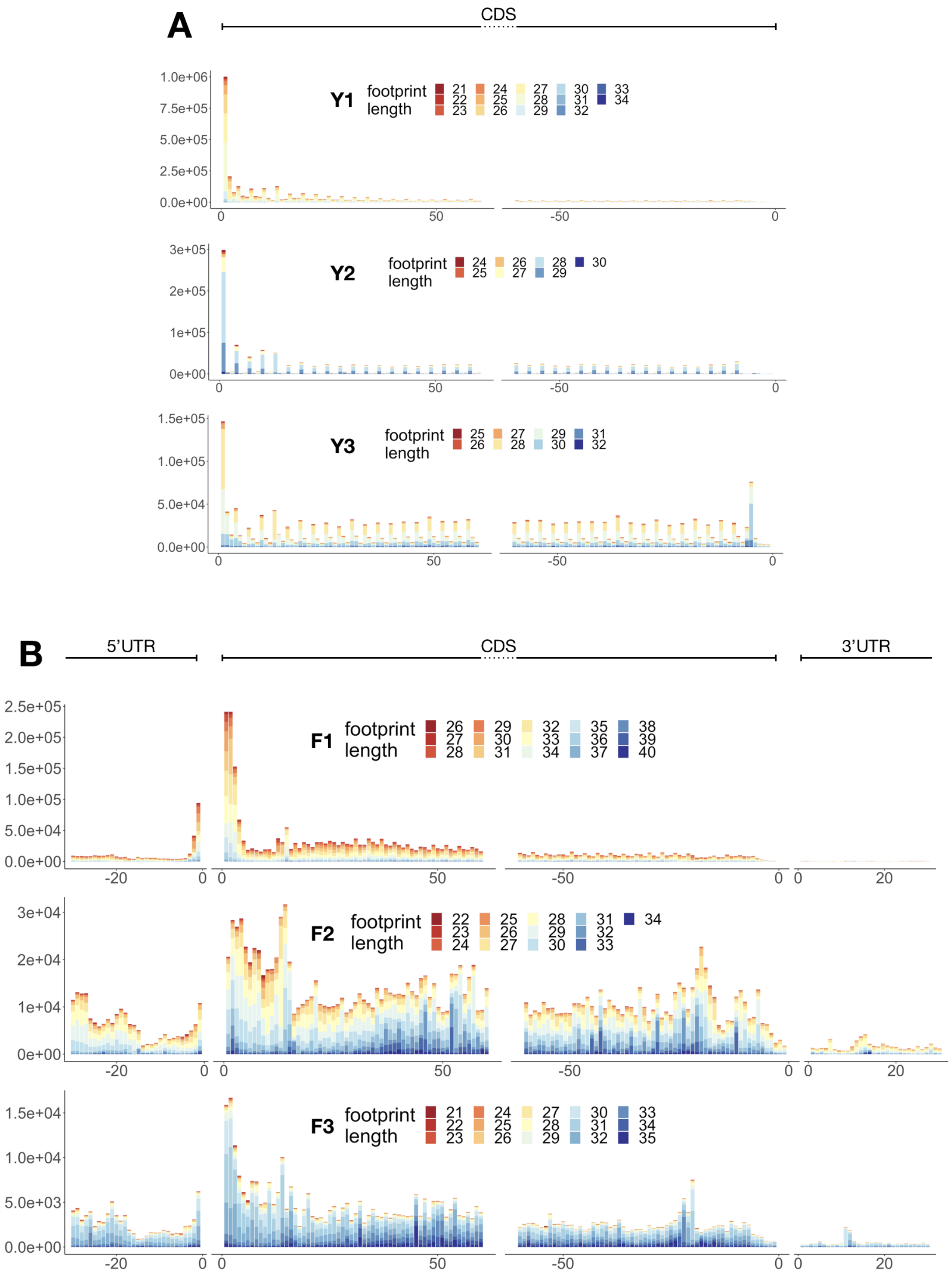
A
CDS
Y1
footprint length
21 22 23 24 25 26 27 28 29 30 31 32 33 34
1.0e+06
7.5e+05
5.0e+05
2.5e+05
0.0e+00
0
50
-50
0
Y2
footprint length
24 25 26 27 28 29 30
3e+05
2e+05
1e+05
0e+00
Y3
footprint length
25 26 27 28 29 30 31 32
1.5e+05
1.0e+05
5.0e+04
0.0e+00
B
5'UTR
CDS
3'UTR
F1
footprint length
26 27 28 29 30 31 32 33 34 35 36 37 38 39 40
2.5e+05
2.0e+05
1.5e+05
1.0e+05
5.0e+04
0.0e+00
-20
20
F2
footprint length
22 23 24 25 26 27 28 29 30 31 32 33 34
3e+04
2e+04
1e+04
0e+00
F3
footprint length
21 22 23 24 25 26 27 28 29 30 31 32 33 34 35
1.5e+04
1.0e+04
5.0e+03
0.0e+00

**Supplementary Figure 1: Ribosome meta-profiles of reads assigned to P-site for each library.** (**A**) yeast, (**B**) fruit fly, (**C**) zebrafish, (**D**) mouse and (**E**) human. On x-axis: last 30 bases of 5′UTR, first and last 60 bases of CDS and first 30 bases of 3′UTR. Ribosome footprints are separated by length, ranging from short (red) to long (blue) fragments. The peaks visible over start codon and the last sense codon (positions 0 to 2 and -5 to -3 on CDS), clear periodicity and high coverage within CDS compared to UTRs indicate proper assignment of reads to P-site. Peaks at start codon and increased coverage on the first few codons of CDSs are clear in CHX and other translation inhibitors-treated libraries (Y1, Y2, F1, F2, F3, Z3, Z5, Z7, M1, M3, H1, H3, H4), while the peak at translation stop is visible in flash-frozen libraries (Y3, Z1, Z2, Z4, Z6, M2, H2).

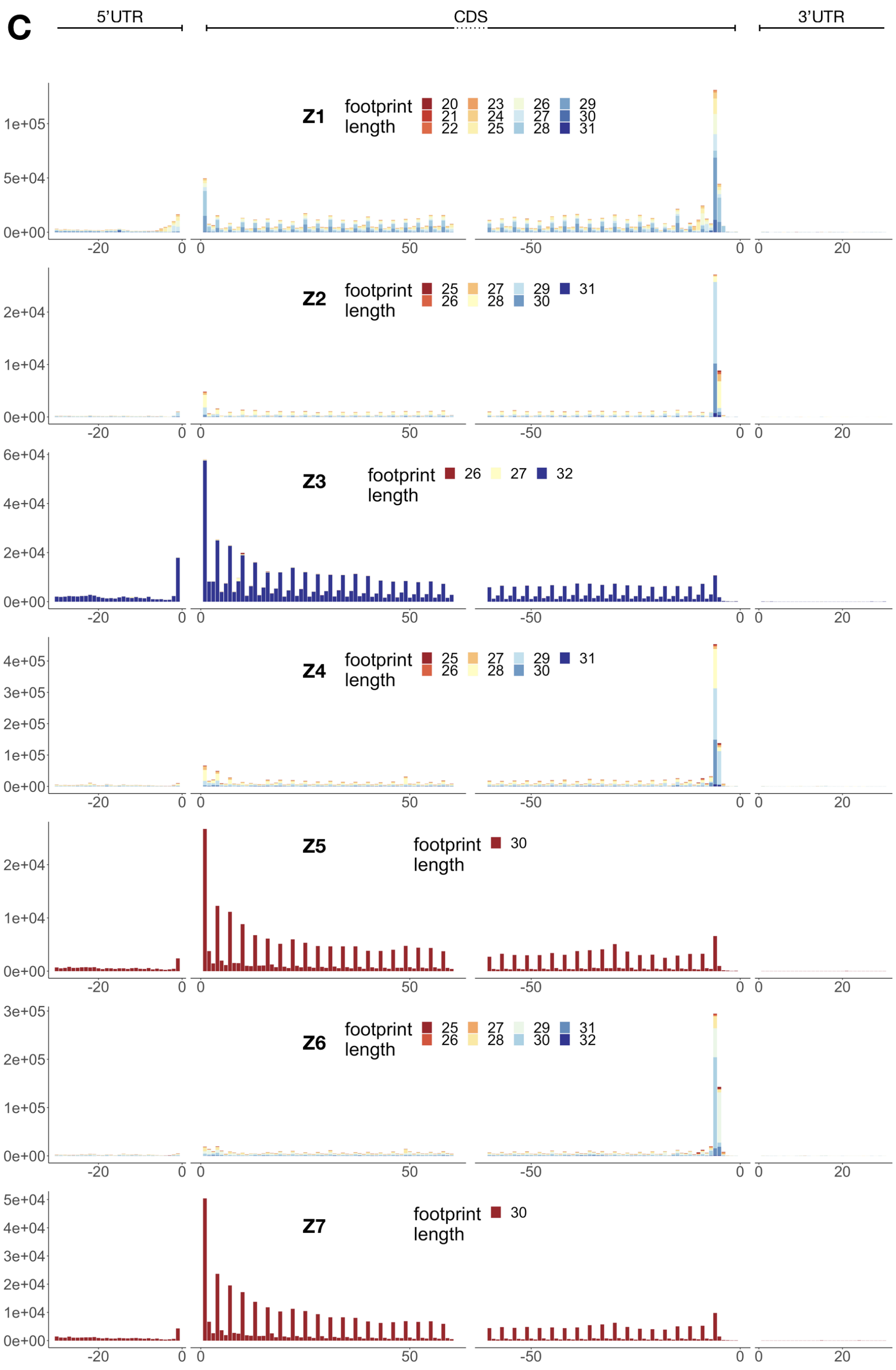


**Supplementary Figure 1:** - continuation.

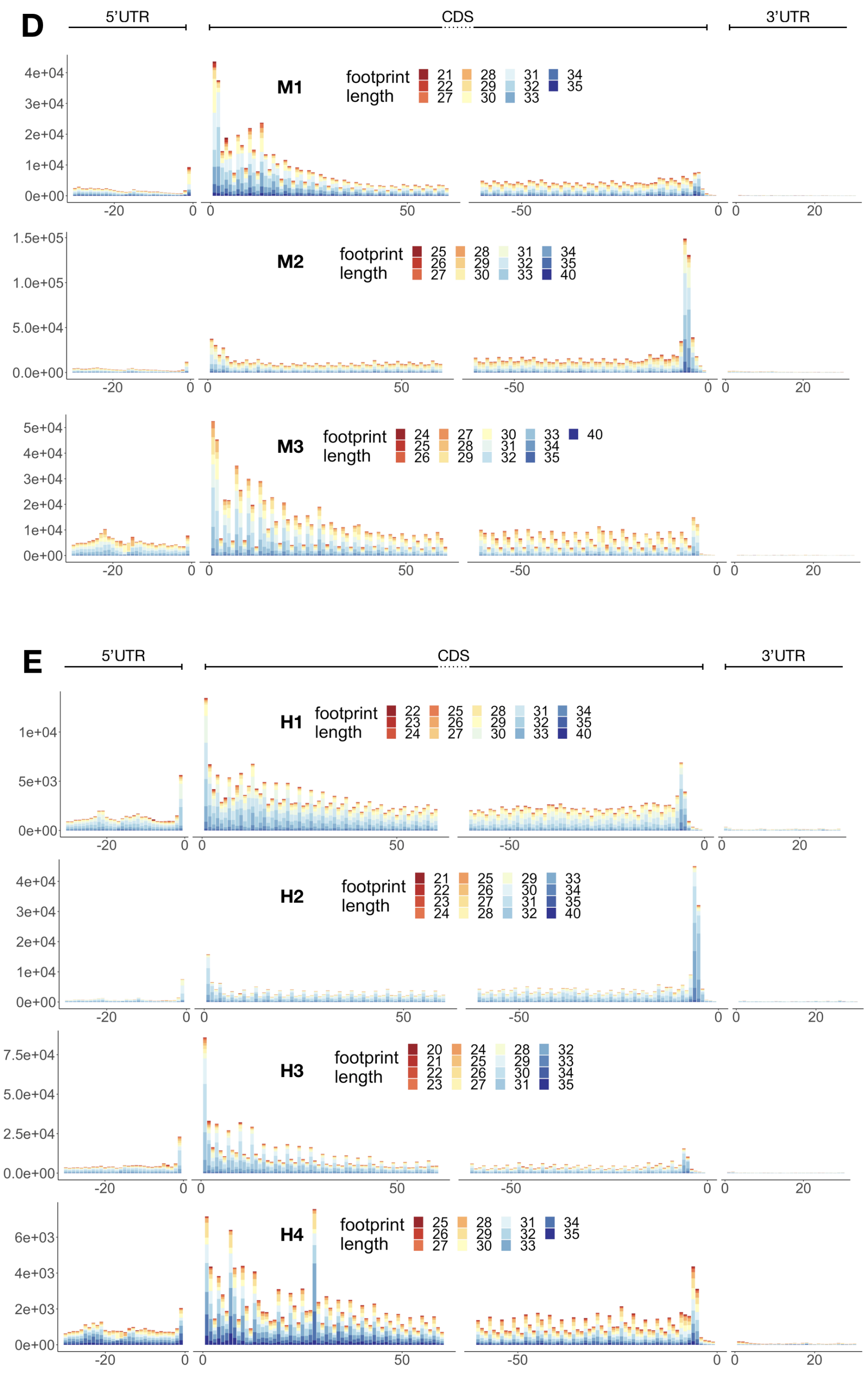


**Supplementary Figure 1:** - continuation.

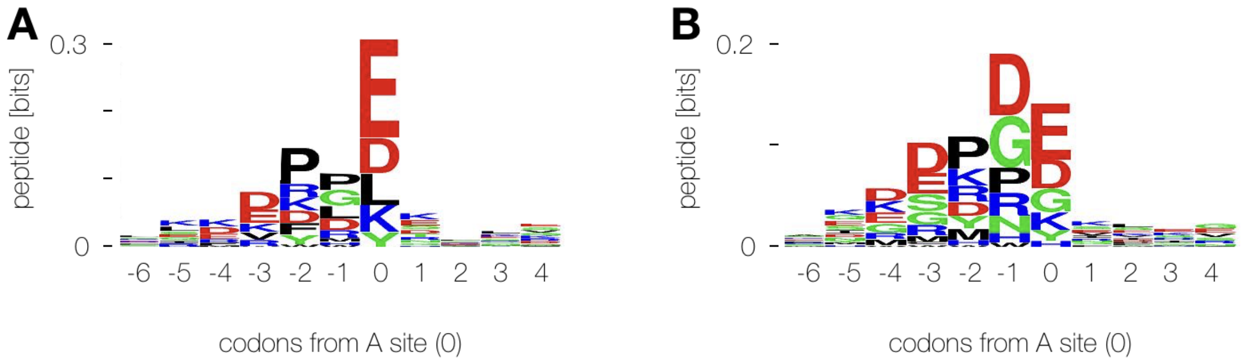


**Supplementary Figure 2: Peptide motif around stall sites.** (**A**) Motif in M2 library, as reported previously in [29] and (**B**) consensus peaks from M1 and M2 libraries.

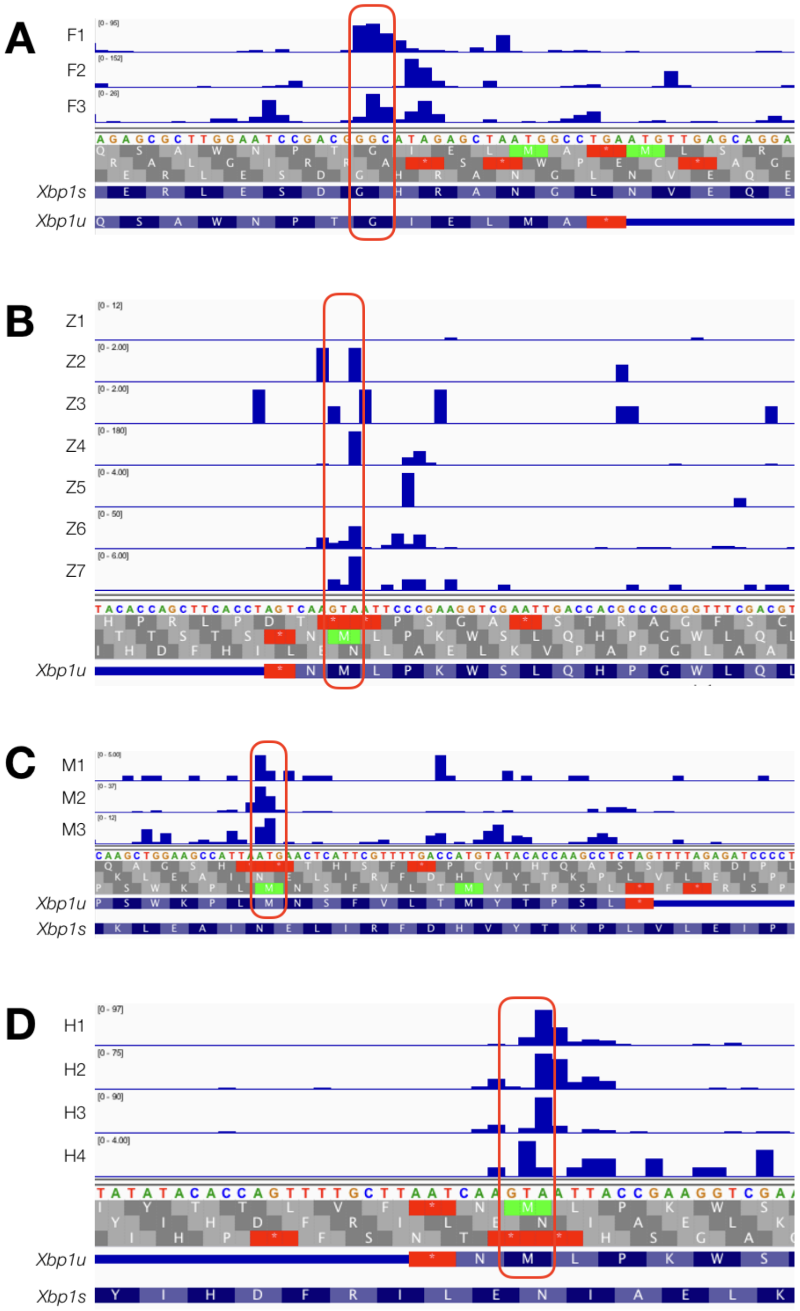


**Supplementary Figure 3: *Xbp1* stall site.** (**A**) Ribosome coverage in fruit fly, (**B**) zebrafish, (**C**) mouse and (**D**) human libraries.

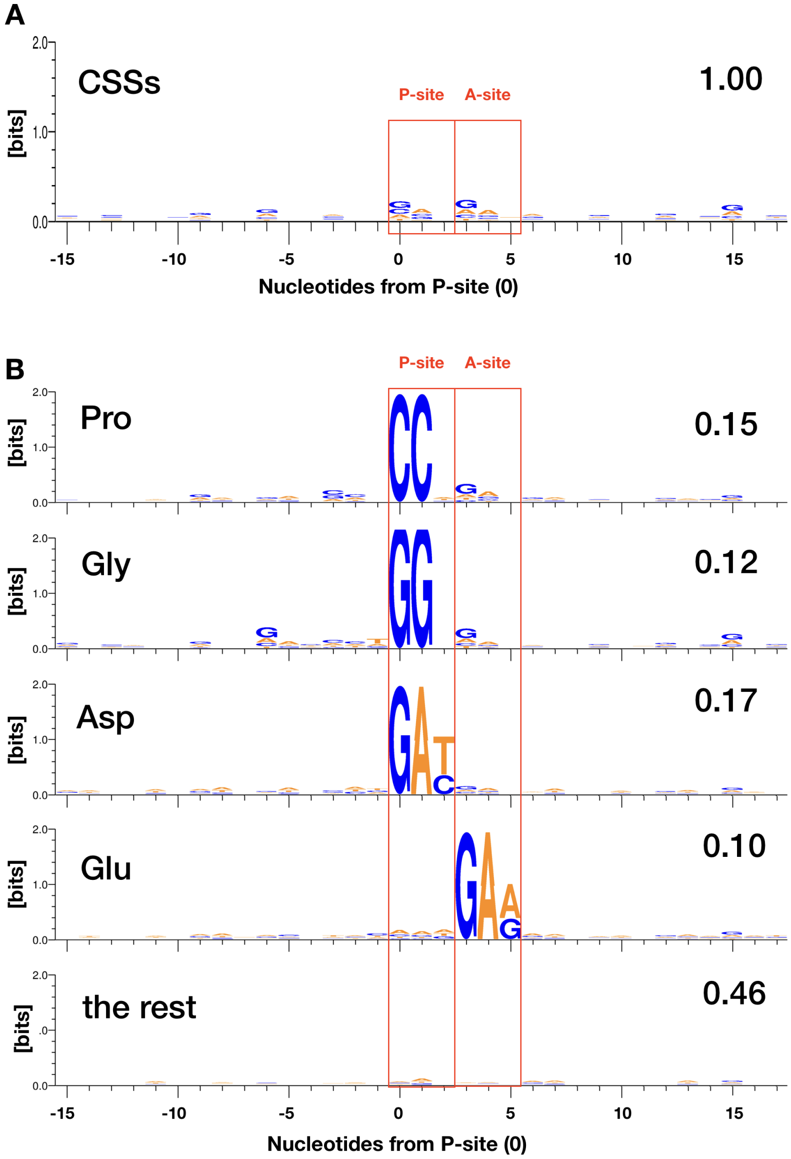


**Supplementary Figure 4: Nucleotide bias around conserved stall sites.** (**A**) Sequence logo for CSSs. (**B**) Sequence logos split by the most significant contributors: prolines, glycines, aspartate at P-site and glutamate at A-site, accounting for 54% CSSs together, and logo of the remaining 46% sequences. The amino acids show biased context, which is not visible on control logos for all of these amino acids extracted from the same transcripts (not shown).

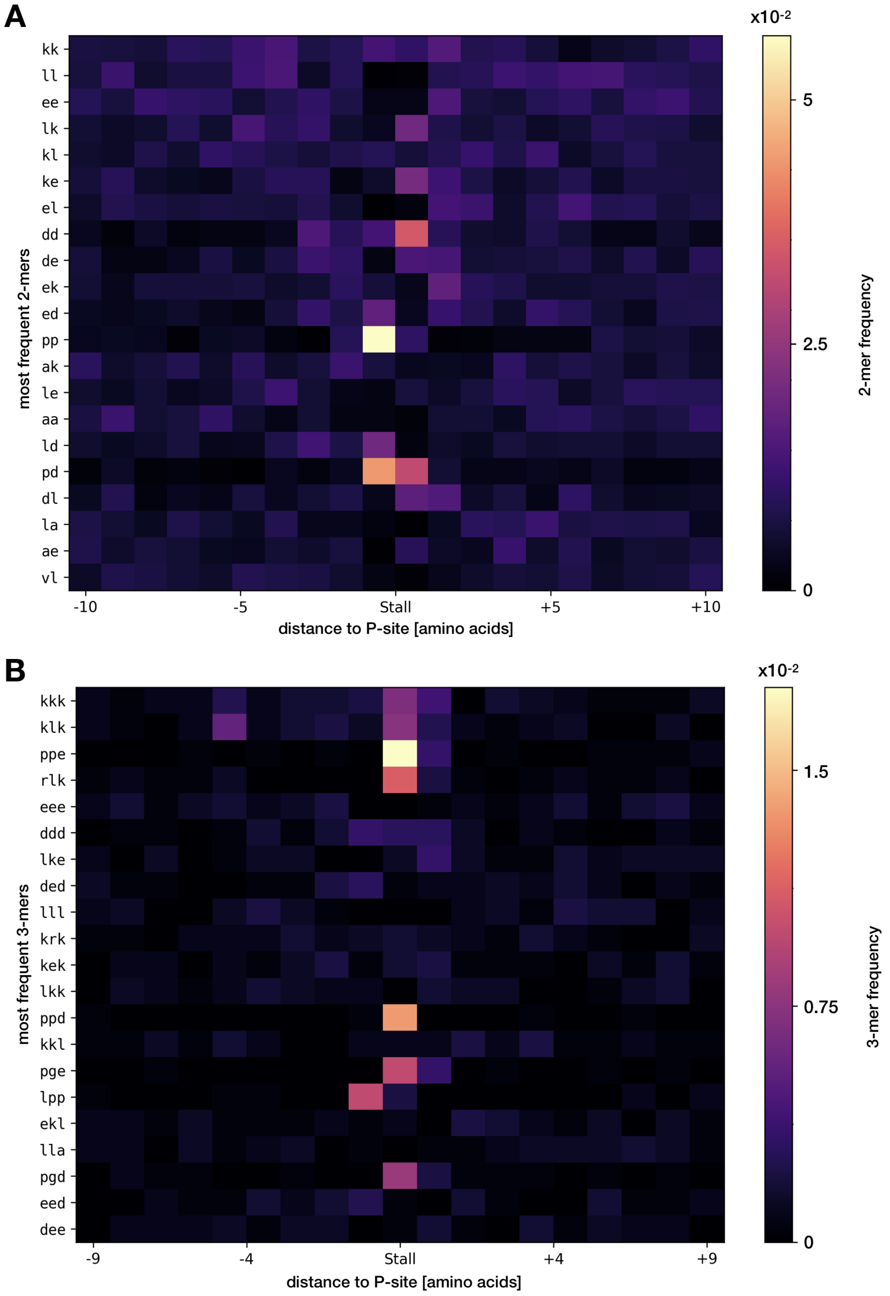


**Supplementary Figure 5: Overrepresentation of amino acid k-mers around stall sites.** (**A**) 2-mers and (**B**) 3-mers, sorted by frequency of occurrence in CSSs, top to bottom.

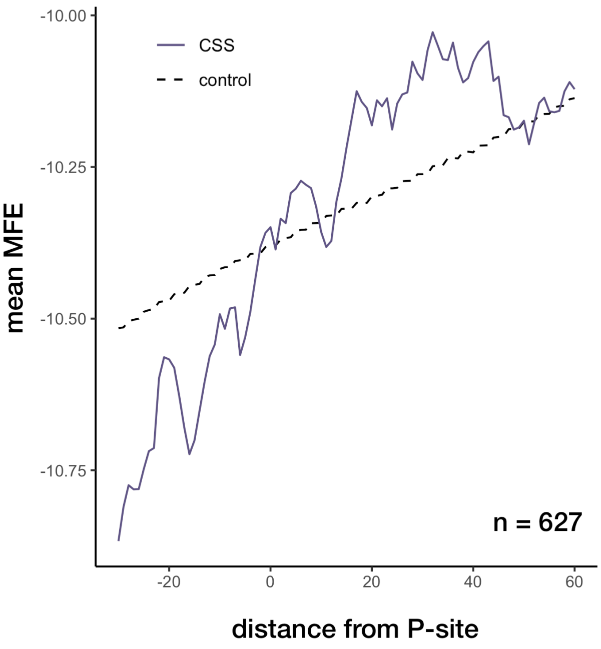


**Supplementary Figure 6: Structure around stall sites.** Mean minimum free energy of regions around CSSs that were not explained by sequence features, compared to background of random sites on the same transcripts.

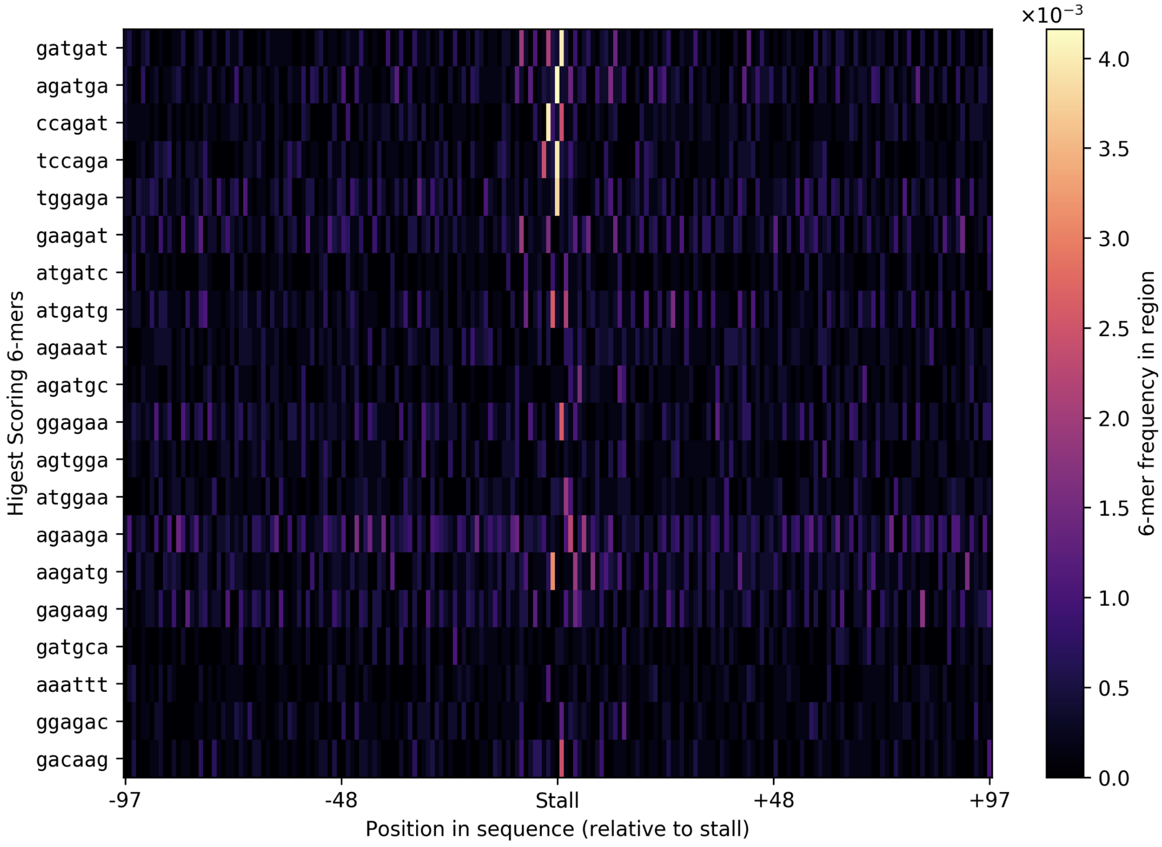


**Supplementary Figure 7: Nucleotide 6-mers flanking CSSs.**

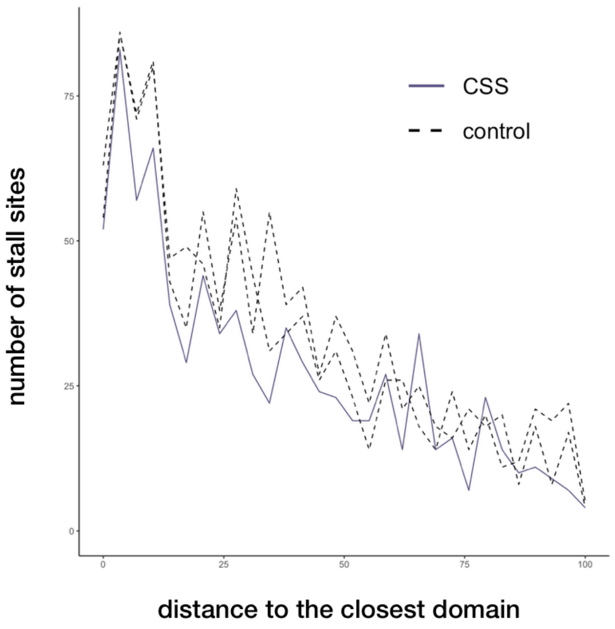


**Supplementary Figure 8: Distance to transmembrane domains.** Distance of the CSSs to the end of the closest upstream protein domain, compared to random controls.

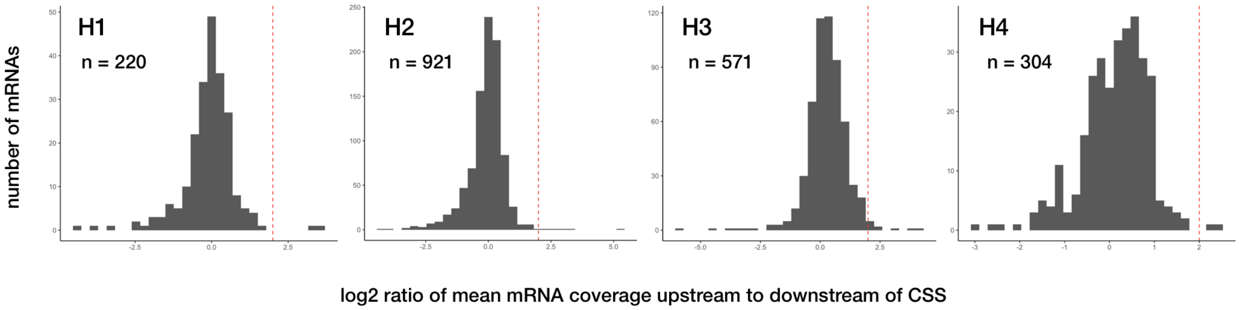


**Supplementary Figure 9: Stalling and degradation.** Change fold of the mRNA coverage upstream of the CSS to downstream for the four human libraries. Red dotted line is set at log2 ratio = 2, with transcripts to the right being possible candidates for stalling-regulated mRNA degradation.

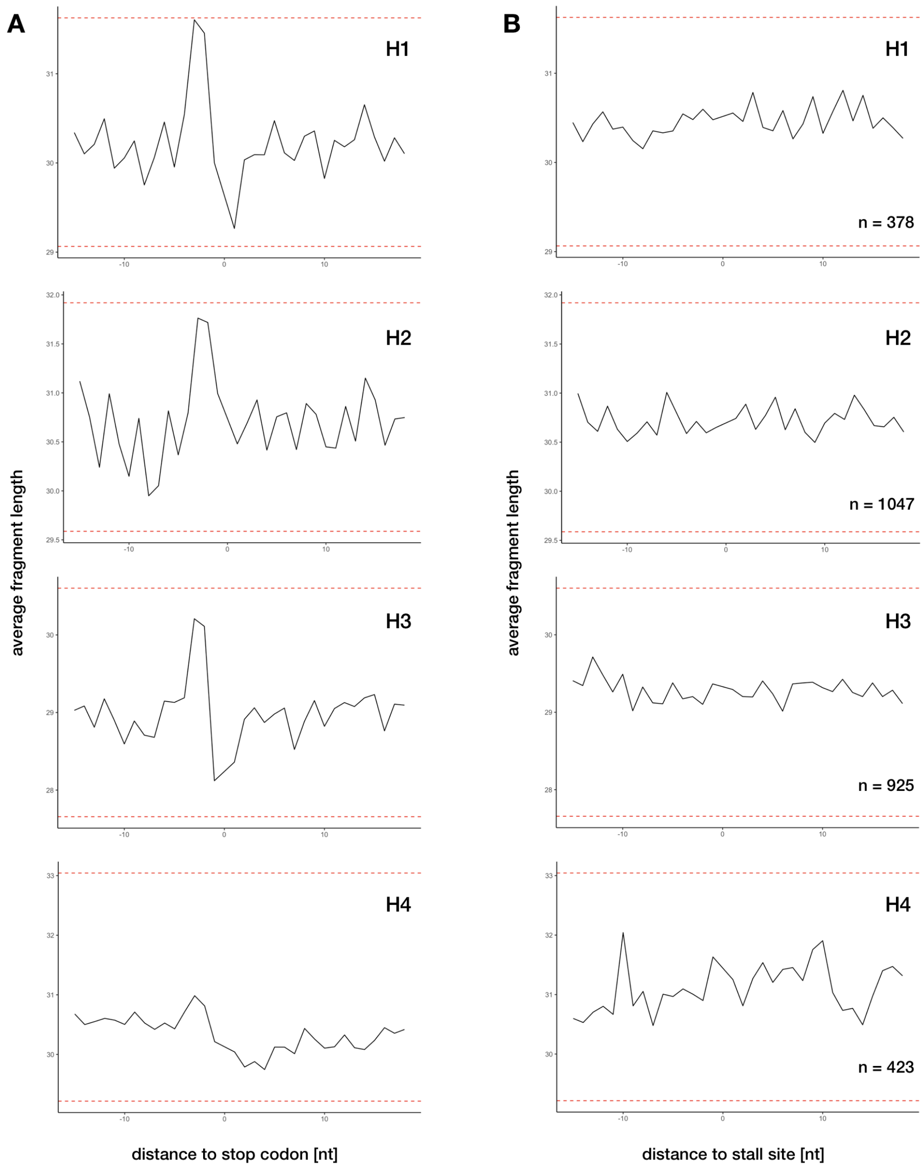


**Supplementary Figure 10: Shift in fragment length distribution.** Average fragment length distribution around (**A**) stop codons and (**B**) CSSs present in four human libraries, H1-H4. The red lines indicate 10% tails of the fragment length distribution calculated from the whole libraries.
